# ModelArray: a memory-efficient R package for statistical analysis of fixel data

**DOI:** 10.1101/2022.07.12.499631

**Authors:** Chenying Zhao, Tinashe M. Tapera, Joëlle Bagautdinova, Josiane Bourque, Sydney Covitz, Raquel E. Gur, Ruben C. Gur, Bart Larsen, Kahini Mehta, Steven L. Meisler, Kristin Murtha, John Muschelli, David R. Roalf, Valerie J. Sydnor, Alessandra M. Valcarcel, Russell T. Shinohara, Matthew Cieslak, Theodore D. Satterthwaite

**Affiliations:** Lifespan Informatics and Neuroimaging Center (PennLINC), Department of Psychiatry, Perelman School of Medicine, University of Pennsylvania, Philadelphia, PA, 19104, USA; Penn/CHOP Lifespan Brain Institute, Perelman School of Medicine, Children’s Hospital of Philadelphia Research Institute, Philadelphia, PA 19104, USA; Department of Bioengineering, School of Engineering and Applied Science, University of Pennsylvania, Philadelphia, PA 19104, USA; Department of Psychiatry, Perelman School of Medicine, University of Pennsylvania, Philadelphia, PA 19104, USA; Program in Speech and Hearing Bioscience and Technology, Harvard University, Cambridge, MA 02139, USA; Department of Biostatistics, Johns Hopkins Bloomberg School of Public Health, Baltimore, MD 21205, USA; Penn Statistics in Imaging and Visualization Center, Department of Biostatistics, Epidemiology and Informatics, University of Pennsylvania, Philadelphia, PA 19104, USA; Center for Biomedical Image Computation and Analytics, University of Pennsylvania, Philadelphia, PA 19104, USA

**Keywords:** Fixel-based analysis, statistical analysis, software, development, big data, MRI

## Abstract

Diffusion MRI is the dominant non-invasive imaging method used to characterize white matter organization in health and disease. Increasingly, fiber-specific properties within a voxel are analyzed using fixels. While tools for conducting statistical analyses of fixel data exist, currently available tools are memory intensive, difficult to scale to large datasets, and support only a limited number of statistical models. Here we introduce ModelArray, a memory-efficient R package for mass-univariate statistical analysis of fixel data. With only several lines of code, even large fixel datasets can be analyzed using a standard personal computer. At present, ModelArray supports linear models as well as generalized additive models (GAMs), which are particularly useful for studying nonlinear effects in lifespan data. Detailed memory profiling revealed that ModelArray required only limited memory even for large datasets. As an example, we applied ModelArray to fixel data derived from diffusion images acquired as part of the Philadelphia Neurodevelopmental Cohort (n=938). ModelArray required far less memory than existing tools and revealed anticipated nonlinear developmental effects in white matter. Moving forward, ModelArray is supported by an open-source software development model that can incorporate additional statistical models and other imaging data types. Taken together, ModelArray provides an efficient and flexible platform for statistical analysis of fixel data.

**HIGHLIGHTS:**

- ModelArray is an R package for mass-univariate statistical analysis of fixel data
- ModelArray is memory-efficient even for large-scale datasets
- ModelArray supports linear and nonlinear modeling and is extensible to more models
- ModelArray facilitates easy statistical analysis of large-scale fixel data

**Graphical abstract:** 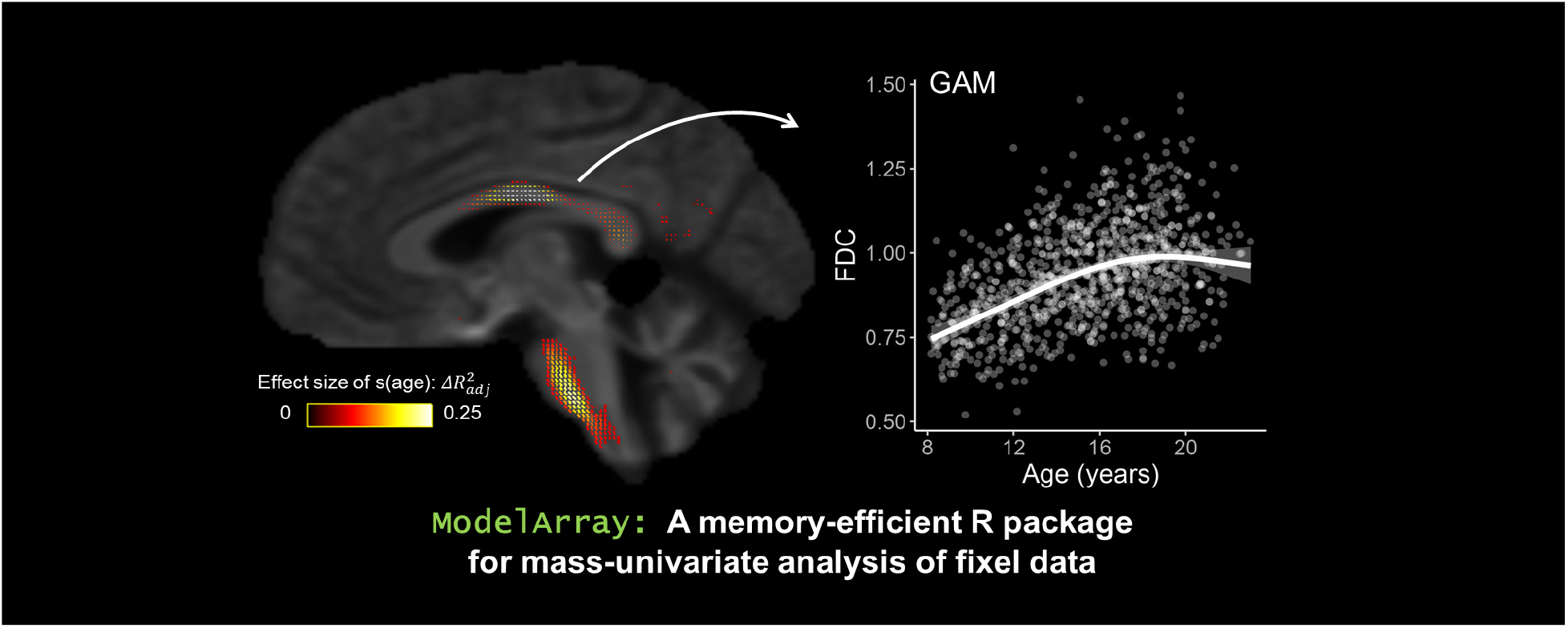

## INTRODUCTION

Diffusion MRI (dMRI) is the dominant method used to non-invasively study white matter organization in the human brain. The most commonly used method for modeling the diffusion signal is diffusion tensor imaging (DTI; Basser & Pierpaoli, 1996). However, DTI cannot effectively model two or more crossing fibers within a given voxel; crossing fibers are thought to comprise up to ~90% of white matter (WM) voxels (Jeurissen et al., 2013; Schilling et al., 2018; Yeh et al., 2013). One method for addressing crossing fibers that is increasingly ascendant is fixel-based analysis (FBA; Raffelt et al., 2015, 2017). A *fixel* refers to a specific fiber population in a voxel; with FBA, multiple distinct fiber populations can be estimated within a voxel and multiple fiber-specific properties can be quantified (Raffelt et al., 2015, 2017). The FBA pipeline typically includes two parts. First, fixel data is generated for each participant in a sample and quantified according to standard measures like fiber density (FD), fiber-bundle cross-section (FC), or their combination – fiber density and cross-section (FDC). Second, the high-dimensional fixel data from a sample is often analyzed in template space using mass-univariate hypothesis testing; this often relies upon connectivity-based fixel enhancement (CFE) as implemented in MRtrix (https://www.mrtrix.org/; Tournier et al., 2019).

However, current tools have two limitations. First, CFE has high memory demands, which may scale by image resolution and sample size (Raffelt et al., 2015). This impedes the application of FBA in large-scale dMRI data resources that include thousands of participants; e.g., the Philadelphia Neurodevelopmental Cohort (PNC; Satterthwaite et al., 2014), the Human Connectome Project (HCP; Van Essen et al., 2013), or the Healthy Brain Network (HBN; Alexander et al., 2017). When faced with such large data resources, investigators often opt to reduce the dimensionality of the data and use regional summary measures, even if it is not scientifically optimal.

Second, the statistical models supported by MRtrix for FBA are currently limited to the general linear model (GLM). This may not be optimal for lifespan studies where effects of interest are often nonlinear (e.g., Bethlehem et al., 2022; Lebel et al., 2012). Ideally, a statistical analysis toolset should be extensible to incorporate diverse statistical models. R (https://www.R-project.org; R Core Team, 2021) is a popular open-source statistical programming software, and it supports a myriad of statistical functionality. Generalized additive models (GAMs; Wood, 2001, 2004) are among the most widely used approaches to model nonlinear effects of interest in R. GAMs can rigorously model both linear and nonlinear effects by applying a penalty that helps avoid over-fitting; this approach is particularly valuable in high-dimensional data settings – cases when hundreds of thousands of fixels are present – where it is difficult to conduct detailed model diagnostics.

To address these limitations, we introduce ModelArray (https://pennlinc.github.io/ModelArray/), a memory-efficient R package for statistical analysis of fixel data. To maximize memory efficiency, ModelArray does not load the entire fixel data into the memory. Instead, it only reads a limited block of data when needed by leveraging the Hierarchical Data Format 5 (HDF5) file format and DelayedArray package in R (Pagès et al., 2021), At present, ModelArray supports linear models and GAMs, but it is by design extensible and can incorporate many statistical models implemented in R. To demonstrate ModelArray’s scalability, functionality, and extensibility, we profiled its memory usage and applied it to examine nonlinear patterns of brain development using fixel data from the PNC (n = 938). As described below, ModelArray allows for efficient and flexible analysis of fixel data in large scale data resources.

## MATERIALS AND METHODS

### Overview

ModelArray is an R package for mass-univariate hypothesis testing of fixel data that is designed to be scalable for large datasets. We chose R as the platform as it is among the most widely used platforms for statistical computing. This feature facilitates the potential to easily incorporate diverse statistical models. ModelArray takes the fixel-wise data as input, after it has been converted to the HDF5 format by its companion software ConFixel (https://github.com/PennLINC/ConFixel). Fixel-wise data with metrics such as FD, FC, and FDC can be calculated in existing software such as MRtrix (Tournier et al., 2019). ModelArray performs statistical analysis for each fixel based on the statistical formula a user provides, and finally saves statistical output as images via ConFixel. These output images can then be viewed in widely-used visualization tools such as MRView from MRtrix (https://www.mrtrix.org/; Tournier et al., 2019).

### Software design and memory efficiency

We capitalized upon the R package DelayedArray (Pagès et al., 2021) to maximize memory efficiency. Of note, the term “memory” is used in this paper to refer to the computer’s memory (RAM) used by software (including data loaded into the memory), and “disk” or “disk space” refers to the hard disk space where the files (e.g., an HDF5 file) are stored. ModelArray wraps fixel data on disk into a DelayedArray object, allowing common array operations such as indexing (e.g., extracting values of a specific fixel from a matrix) or transposing to be performed without loading the on-disk object into memory. DelayedArray objects store their component data in an HDF5 file, and operations on a DelayedArray object are executed in a memory-efficient, “delayed” way (where most R operations are processed on-demand and *en masse*). The result is a memory-efficient and easy-to-use R interface for a large and hierarchical on-disk dataset. After being generated by ConFixel (see below), an HDF5 file of fixel data contains a scalar matrix (fixels by participants), basic information of fixels and voxels (e.g., lookup tables of the directions of fixels and the coordinates of voxels that contain fixels), and, once calculated by ModelArray, one or more result matrices (fixels by statistical metrics). Leveraging DelayedArray, HDF5 format, and the supporting R package HDF5Array (Pagès, 2021), the on-disk fixel data can be accessed and manipulated while minimizing memory requirements.

### ModelArray workflow

ModelArray is packaged with the companion software ConFixel for converting fixel data to the expected file format (see **Figure 1**). Specifically, ConFixel is a Python-based command-line interface software, and it converts between the original MRtrix image format (.mif) and the HDF5 file format (.h5) used for ModelArray. After the file format conversion, ModelArray generates a ModelArray-class object for representing the on-disk HDF5 file. ModelArray uses the S4 Object Oriented Programming (OOP) model which gives users easy access to the scalar matrix, the source .mif file list, one or more results matrices (if any), and the file path to the HDF5 file. When fitting models, ModelArray iterates across all fixels in the scalar matrix but only reads a limited block of data for each current fixel in order to reduce memory usage. For each fixel, the software fits a model for the participant-level phenotypes of interest – such as age, sex, or diagnosis, which are loaded from a separate CSV file provided by the user – and generates the statistical outputs for each fixel, such as *p*-values, coefficient estimations, and *t*-statistics. After generating the result matrix of fixel-wise statistics, ModelArray will calculate corrected *p*-values using the False Discovery Rate (FDR) and export the final result matrix back into the input HDF5 file. Finally, ConFixel converts the HDF5 file’s results matrix into a list of .mif files that are readable by widely-used visualization tools such as MRView from MRtrix (https://www.mrtrix.org/; Tournier et al., 2019).

**Figure 1.**
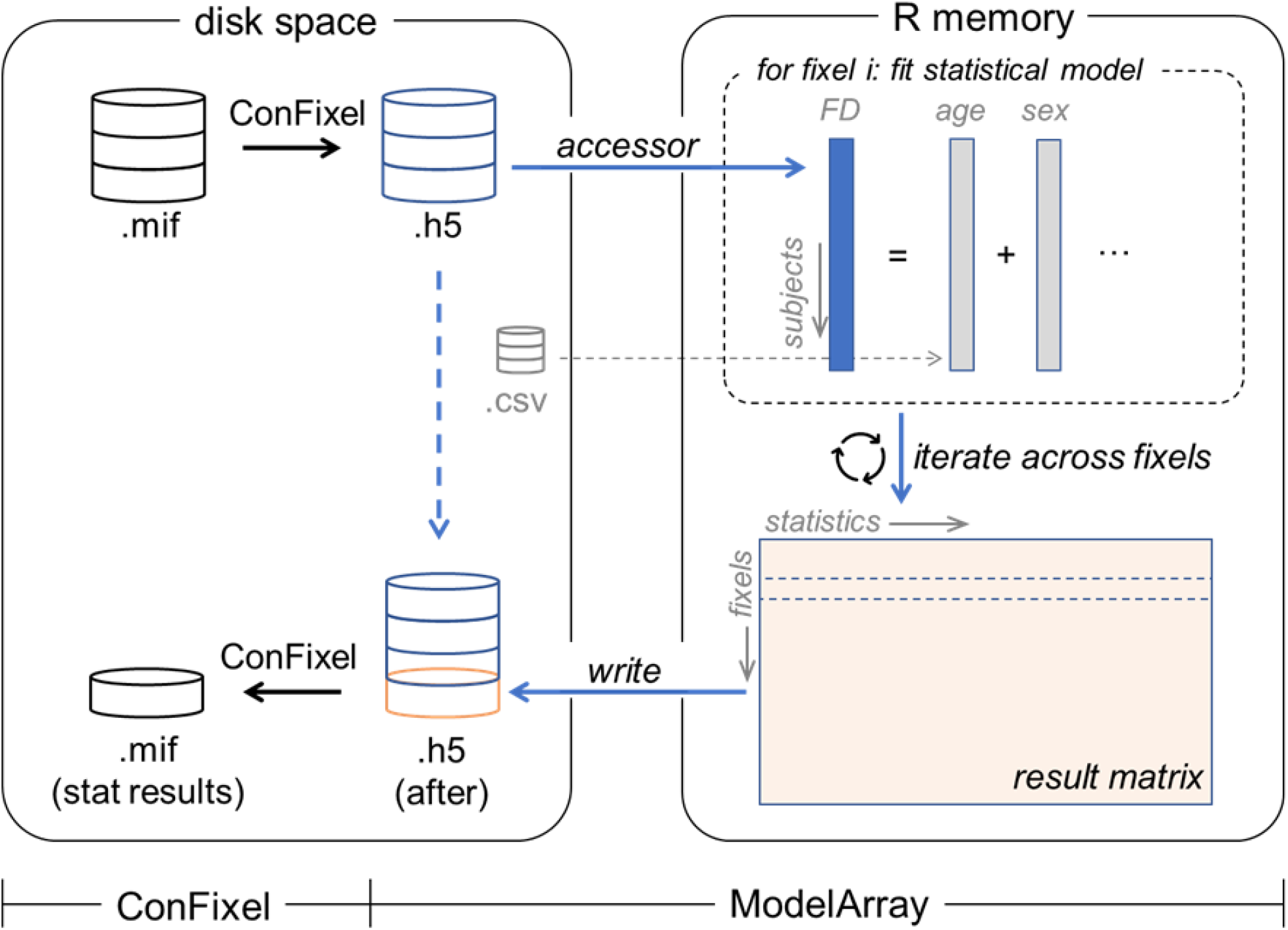
Schematic of ModelArray and its companion converter ConFixel. The original fixel data (.mif files) are first converted into an HDF5 file (.h5) using ConFixel (top of the left box). ModelArray provides easy access to fixel data in the HDF5 file (“accessor”). When performing statistical analysis of each fixel (top of the right box), to reduce memory usage, only a limited block of fixel data is read into the memory. Using the phenotypes of interest (e.g.,: age, sex; provided by a CSV file), ModelArray fits a statistical model and calculates statistical output for each fixel. After iterating across fixels, the result matrix is generated (bottom of the right box) and saved to the original HDF5 file on disk by ModelArray (“write”). Finally, ConFixel converts the result matrix in this HDF5 file into a list of .mif files ready to be viewed (bottom of the left box).

### ModelArray functions

ModelArray provides functions for model fitting and writing statistical results. At present, ModelArray supports linear models (`ModelArray.lm()`) as well as GAMs with and without penalized splines (`ModelArray.gam()`). Model fitting can be accelerated by requesting more CPU cores for parallel computing. ModelArray writes the rich statistical output of R into an HDF5 file using the `writeResults()` function. This HDF5 file is then converted to a list of .mif files with ConFixel for viewing, as described above. Default statistical output from ModelArray includes several maps for each model term (e.g., coefficient, *t*-statistic, raw and FDR-corrected *p*-values), as well as maps regarding the overall model fit (e.g., adjusted *R*-squared, raw and FDR-corrected *p*-values from the model *F*-test in linear models). New statistical models can be easily added by any GitHub contributor following the same workflow as existing ones (`ModelArray.lm()` and `ModelArray.gam()`); see developer documentation at: https://pennlinc.github.io/ModelArray/articles/doc_for_developer.html. Thus, ModelArray is extensible to many diverse statistical methods used in R.

### Evaluation data

To evaluate ModelArray, we used the fixel data generated from the Philadelphia Neurodevelopmental Cohort (PNC; Satterthwaite et al., 2014). Here we provide a brief summary of the data and methods including participant inclusion, image acquisition, image quality assurance, diffusion MRI preprocessing, and fixel-based analysis. In total, we included n=938 participants (521 female, 417 male) aged 8-23 years old. Participants were excluded due to lack of diffusion imaging data, abnormalities in brain structure, major health conditions, missing B0 field map, poor image quality, etc. All the dMRI data underwent a rigorous manual and automated quality assessment as previously described (Roalf et al., 2016).

MRI scans were acquired on a Siemens TIM Trio 3T scanner. Diffusion MRI scans were acquired with a twice-refocused spin-echo (TRSE) single-shot echo-planar imaging (EPI) sequence. The sequence included 64 diffusion-weighted images of b = 1000 s/mm^2^ as well as 7 interspersed b = 0 images; these images were acquired over two scan runs. The in-plane resolution was 1.875×1.875 mm^2^, slice thickness was 2 mm without gap. In addition, a B0 field map was also acquired for distortion correction of dMRI data. In-scanner motion during the dMRI scan was quantified as the root mean squared displacement (mean relative RMS); this was calculated from 7 b = 0 volumes interspersed over the course of the dMRI scan (Roalf et al., 2016). Motion was included as a covariate when modeling age effects using GAMs (described below). Diffusion images were processed with QSIPrep (https://github.com/PennBBL/qsiprep: Cieslak et al., 2021). This process included denoising, distortion correction, and head motion correction. Finally, the images were resampled to AC-PC alignment with 1.25 mm isotropic voxels.

Following preprocessing, fixel-based analysis was performed using MRtrix (https://www.mrtrix.org/, version v3.0RC3) (Dhollander et al., 2021; Raffelt et al., 2017; Tournier et al., 2019). Briefly, study-specific response functions were calculated using data from 30 representative participants across ages (15M/15F). Fiber orientation distributions (FODs) for all participants were then estimated using single-shell three-tissue constrained spherical deconvolution (CSD) (Tournier et al., 2007). A study-specific FOD template was generated, and participants’ FOD images were registered to this study template. After defining fixels, FDC was quantified and chosen as the metric of interest as it combines both FD and FC and may be more sensitive than FD or FC alone (Dhollander et al., 2021). Finally, the FDC values were smoothed with “connected” nearby fixels to increase the signal-to-noise ratio (Raffelt et al., 2015). To smooth the data, a whole-brain probabilistic tractogram with 2 million streamlines was generated from the FOD template, and a fixel-fixel connectivity matrix based on this tractogram was computed. Lastly, FDC values were smoothed based on this matrix. This procedure yielded fixel data in template space for each participant, which included 602,229 fixels. This fixel data was used by both ModelArray and by the function `fixelcfestats` (Raffelt et al., 2015) in MRtrix for comparison.

### Memory profiling

We evaluated the memory efficiency of ModelArray and compared it to the primary existing tool for fixel-wise statistical analysis: the function `fixelcfestats` in MRtrix (version 3.0.2-193-gdd63cc20) (Raffelt et al., 2015). Memory profiling for both ModelArray and MRtrix was completed using a Linux system by Working Set Size (WSS) Tools for Linux (https://www.brendangregg.com/wss.html). We used a virtual machine on a standalone computer to avoid interference from other users, with memory allocated to the virtual machine = 55 Gigabytes (GB) and total RAM on the computer = 64 GB. Specifically, the resident set size (RSS) –real memory pages currently mapped – was captured by WSS and recorded. We sampled the RSS once every second for both parent and any child processes (if more than one CPU core was used). The total RSS from all processes was calculated by summing the interpolated RSS values at each second, and the maximum RSS used over time was calculated.

To facilitate comparisons in profiling, we used a simple linear model of FDC =intercept + age. To evaluate how memory usage scaled with data size, we examined the full sample (n=938) as well as subsamples of different sizes (n=30, n=100, n=300, n=500, and n=750). Furthermore, memory profiling over different parallelization factors was also performed. During the memory profiling for ModelArray and MRtrix, up to four CPU cores were made available.

We compared memory requirements using both the full dataset (n=938) and smallest subset of data (n=30). For MRtrix `fixelcfestats`, 100 permutations were specified, and *F* tests were performed. In all cases, memory profiling was run three times for each use case, and the median value was reported.

### Application using generalized additive models

The memory benchmarking studies were conducted using linear models, as that functionality is available for both MRtrix and ModelArray. However, in addition, we also demonstrated the use of GAMs in ModelArray for modeling nonlinear developmental effects. Notably, existing tools such as MRtrix only support GLMs and do not easily allow users to model nonlinear developmental effects using GAMs. This application illustrates the extensibility of ModelArray to incorporate diverse statistical models.

For this application, data from all participants (n = 938) was used. Age was modeled as a smooth term s(age) with four basis functions (k=4); sex and in-scanner motion (mean relative RMS displacement) were included as covariates. As in prior work (Pines et al., 2022), the effect size of the age term was quantified as 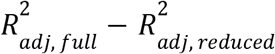, where the 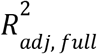 was the adjusted R-squared in the full model, and 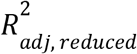 was that in a reduced model that did not
include the age term.

### Open-source software development and release

ModelArray has been developed on GitHub with version controls and all code is openly available on GitHub (see **Data and code availability statements**). Continuous Integration (CI) testing is used to ensure stability and quality assurance. Specifically, we use CircleCI to perform unit tests for all major features of ModelArray. These tests ensure the consistency between the statistical results calculated in ModelArray fitting loop and those calculated in standard R. Once updated code is committed to GitHub, CircleCI automatically builds the software and runs unit tests. If there are any errors, CircleCI will alert the developers to this failure immediately, assuring that updates do not alter software performance.

### Data and code availability statements

ModelArray documentation can be found at https://pennlinc.github.io/ModelArray. All code used to perform memory profiling and application of GAMs is available at https://github.com/PennLINC/ModelArray_paper. The source code for ModelArray is available at https://github.com/PennLINC/ModelArray, and the source code for ConFixel is available at https://github.com/PennLINC/ConFixel. The version of ModelArray used for benchmarking and demonstration was commit SHA-1 0911c4f. The PNC dataset used in this paper is available on dbGAP (https://www.ncbi.nlm.nih.gov/projects/gap/cgi-bin/study.cgi?study_id=phs000607.v3.p2). As part of the software tutorial, example fixel data from 100 PNC participants is openly shared on OSF (https://doi.org/10.17605/OSF.IO/JVEHY).

### Ethics statement

No new data were collected specifically for this paper. The Philadelphia Neurodevelopmental Cohort (PNC; Satterthwaite et al., 2014) was approved by IRBs of the University of Pennsylvania and Children’s Hospital of Philadelphia. All adult participants in the PNC provided informed consent to participate; minors provided assent alongside the informed consent of their parents or guardian.

## RESULTS

### Software walkthrough

Before using ModelArray, two files need to be prepared by the user: an HDF5 (.h5) file of fixel data (example filename here: `example.h5`), and a CSV file of participant’s phenotypes of interest (e.g., age, sex, etc; example filename here: `example.csv`). The HDF5 file can be obtained by applying ConFixel to convert the original fixel data (.mif files) into required HDF5 file format. An example of the usage of ModelArray is displayed in **Figure 2**. After loading the package ModelArray in R (code line #3 in **Figure 2**), a ModelArray-class object `modelarray` was created with the function `ModelArray()`; it represents the fixel data in the HDF5 (.h5) file on disk, including the scalar matrix (fixels by participants) (code line #5). As the entire data was not loaded into memory, this object only required less than 1 Megabytes (MB) for complete n = 938 evaluation data, much less than the HDF5 file size on the disk (2.1 GB). After the data frame of phenotypes was loaded into R (code line #6), mass-univariate analyses using linear models and GAMs were performed with `ModelArray.lm()` and `ModelArray.gam()`, respectively (code line #9-10). The statistical outputs lm.outputs and gam.outputs were saved back to the original HDF5 file with the function `writeResults()` (code line #13-14). These outputs saved in the HDF5 file can be converted back to .mif files by ConFixel for viewing.

**Figure 2.**
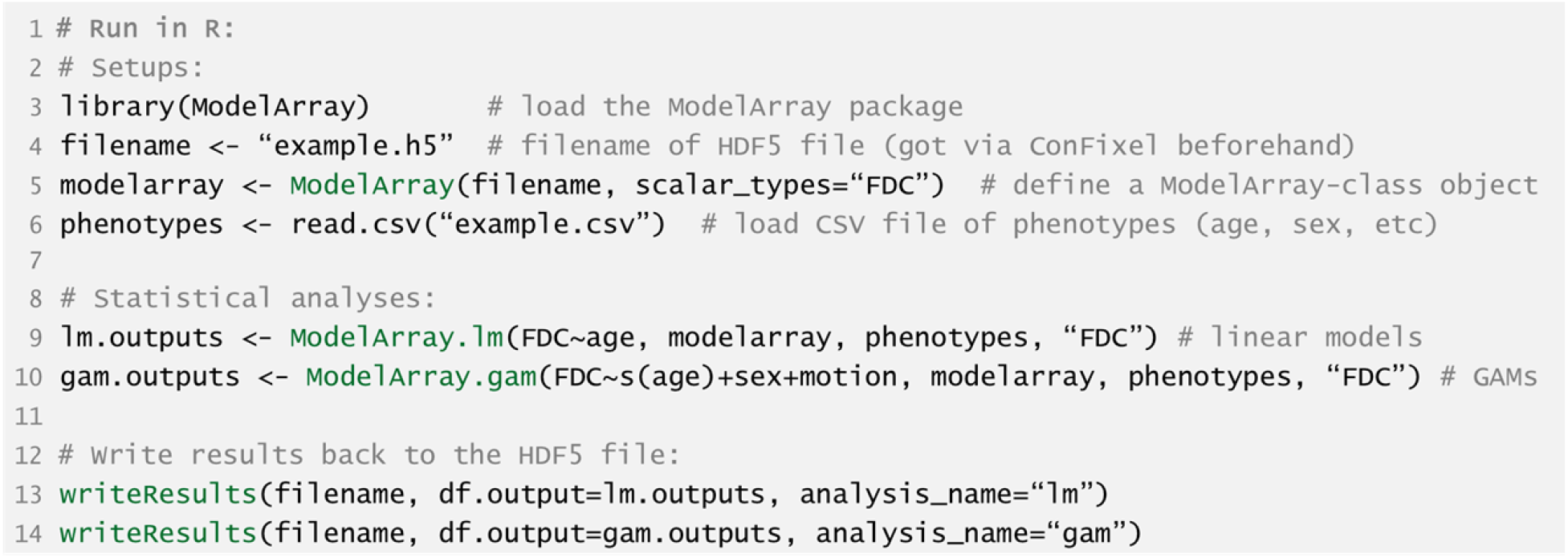
Example R code for executing analysis using ModelArray. ModelArray functions are highlighted in green.

For further details, as part of the comprehensive online documentation, please see the “Walkthrough” of ModelArray and ConFixel (https://pennlinc.github.io/ModelArray/articles/walkthrough.html). This walkthrough can be used in conjunction with openly-shared fixel data from 100 PNC participants, which is available on OSF (https://doi.org/10.17605/OSF.IO/JVEHY).

### ModelArray is memory-efficient and robust to dataset size

We profiled the memory usage of ModelArray and `fixelcfestats` from MRtrix over a range of input data sizes (e.g., number of participants) and parallelization settings. As a first step, we evaluated both the full dataset (n=938) as well as five smaller sub-samples. This initial evaluation was completed using four CPU cores. As the number of participants analyzed increased, ModelArray memory usage only changed minimally (**Figure 3A**). In comparison, MRtrix’s memory requirements scaled with the number of participants included, ultimately requiring 47.79 GB of memory when 938 participants were analyzed (**Figure 3B**).

**Figure 3.**
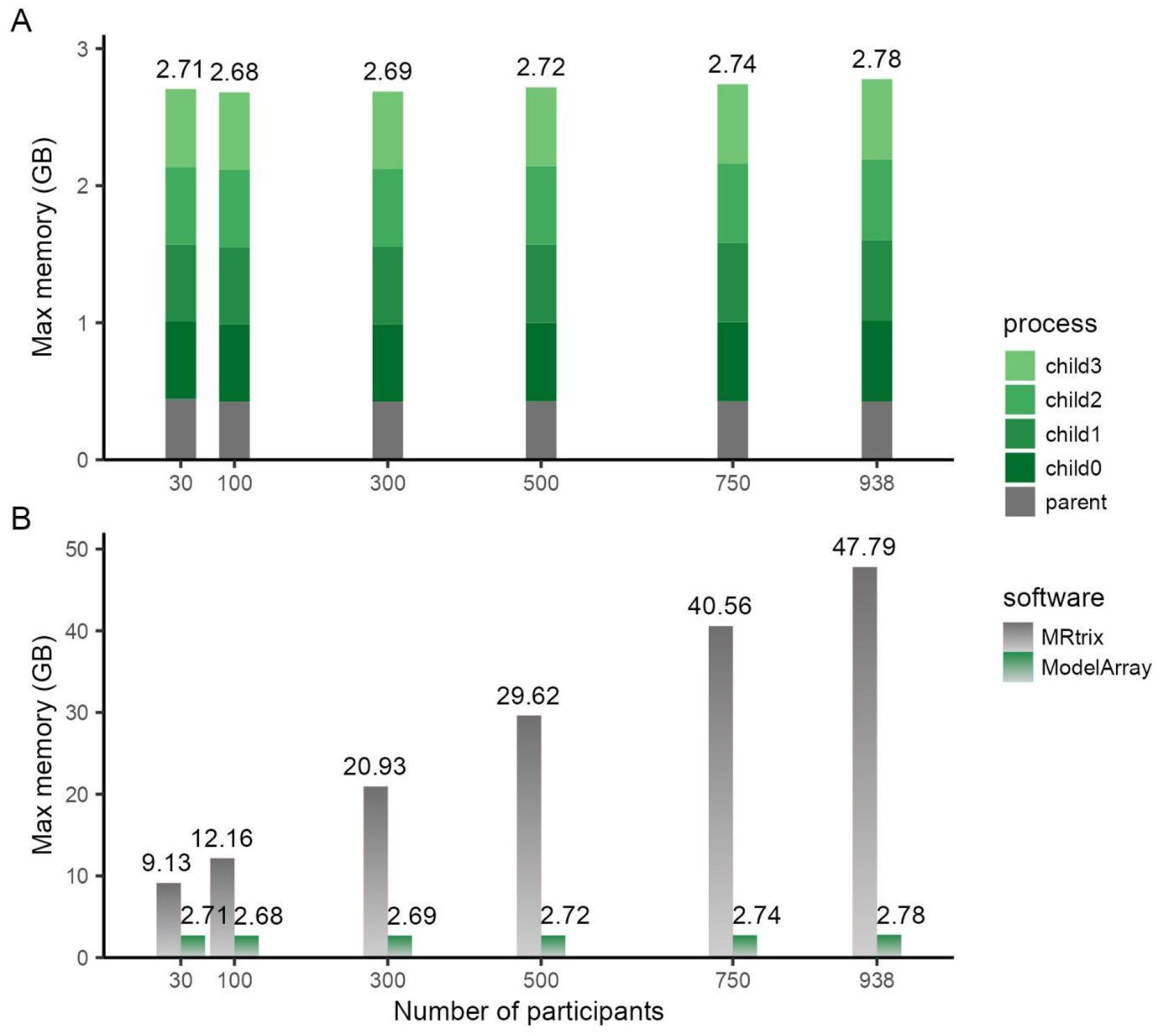
Memory required by ModelArray does not vary by sample size. The maximal memory required by a linear model executed using `ModelArray.lm()` was evaluated when analyzing a range of sample sizes (**A**) and compared with MRtrix (**B**). All models were performed with a parallelization factor of 4.

Next, we examined how parallelization options impacted memory use. As expected, when ModelArray requested more CPUs for analysis of samples of either small (n=30, **Figure 4A**) or large number of participants (n=938, **Figure 4B**), the memory required scaled by the parallelization factor. However, regardless of the parallelization configuration, ModelArray consumed substantially less memory than MRtrix (**Figure 4C** &**D**), especially when analyzing a large number of participants.

**Figure 4.**
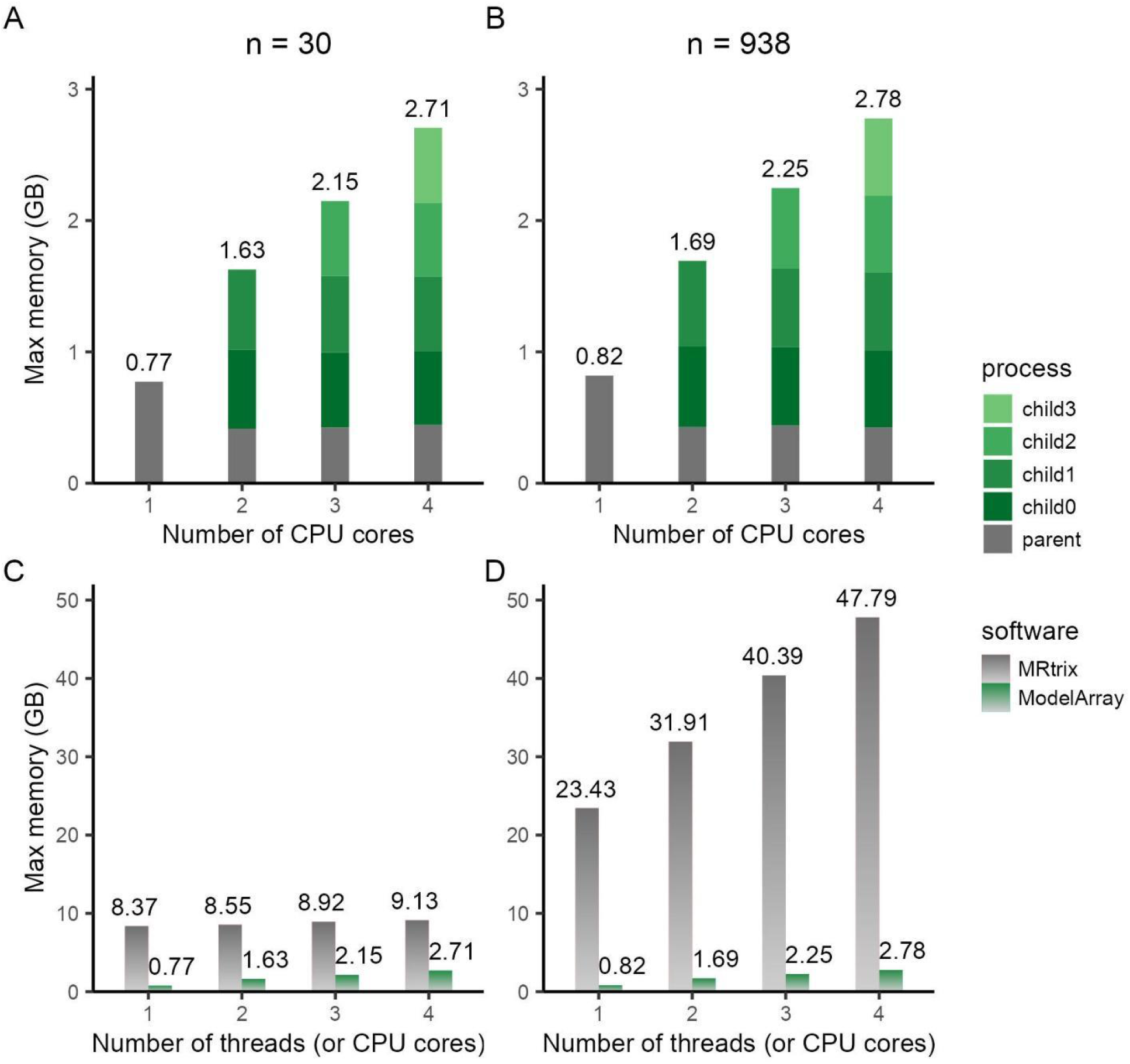
ModelArray is memory-efficient even under different parallelization configurations. Maximal memory usage for a linear model run using `ModelArray.lm()` was evaluated across a sample of n=30 (**A**) and n=938 (**B**) with varying numbers of CPU cores requested (top panels). `ModelArray.lm()` consumed substantially less memory than a comparable analysis using MRtrix in both sample sizes (**C, D**).

### ModelArray captures nonlinear developmental effects

As a final illustration of ModelArray’s functionality and extensibility to diverse statistical models, we also examined nonlinear developmental effects in the PNC using GAMs. Robust nonlinear age effects can be observed in white matter tracts including the corpus callosum (CC) and tracts in the brainstem even at very high statistical thresholds (*p*-value of s(age) < 1×10^-15^, **Figure 5**). To visualize the nonlinear age effects, a cluster in CC was defined with above statistical threshold, and a GAM was fit for FDC averaged in an example 2D slice of this cluster (highlighted in **Figure 5A** by a white arrow). The averaged FDC of these fixels increased throughout childhood and adolescence but then plateaued in young adulthood (**Figure 5B**). The effect size (change in adjusted *R^2^*) of age in this fitted GAM was 0.204.

**Figure 5.**
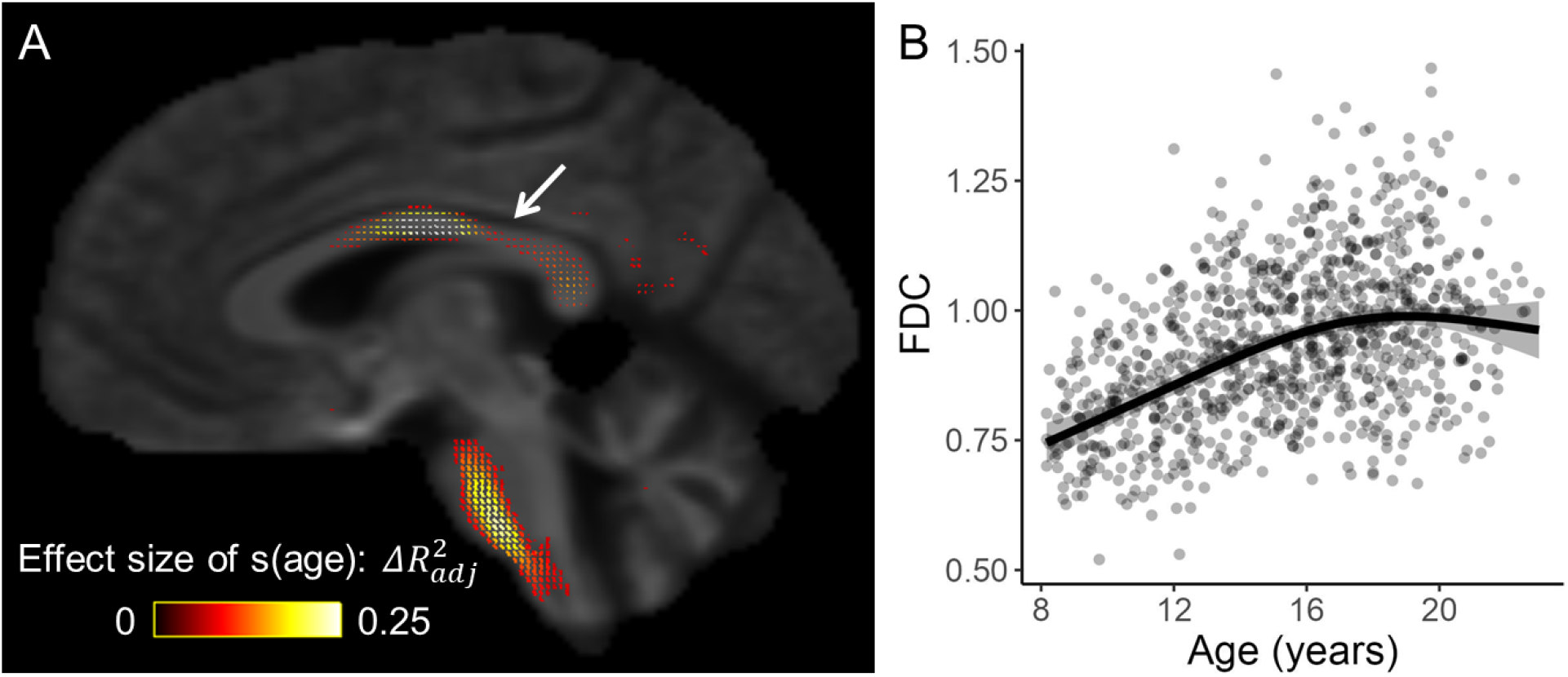
ModelArray allows memory-efficient estimation of nonlinear effects. Fixel-wise GAM fitted with `ModelArray.gam()` revealed nonlinear FDC changes with age in childhood and adolescence (n = 938). The GAM also included sex and motion quantification as covariates. (**A**) Fixels whose FDC was significantly associated with age (*p*-value of s(age) < 1×10^-15^); fixels are colored by effect size of s(age). Background image is the FOD template. (**B**) GAM fit for FDC averaged in the 2D slice of the cluster in CC highlighted in panel **A** by a white arrow.

## DISCUSSION

Despite the advantages of representing diffusion imaging data as fixels, FBA is a relatively new framework compared to voxel-based analysis, and relatively few analytic tools are currently available for statistical analysis of fixel data. ModelArray is an R package for mass-univariate statistical analysis of fixel data. As discussed below, ModelArray allows for both linear and nonlinear modeling of fixel data in large datasets while only requiring modest amounts of memory.

### Scalability to large-scale data resources

Large-scale neuroimaging datasets enhance statistical power and the reliability of findings in studies of individual differences (Marek et al., 2022). However, as data size grows, memory requirements often become quite large when performing group-level statistical analysis. As our benchmarking studies demonstrate, an existing tool for fixel-wise statistical analysis of fixel data (MRtrix) scales with sample size, which can be problematic for with limited computational resources. To address this challenge, we designed ModelArray to minimize memory requirements by only reading data into memory as needed. Our benchmarking studies illustrated that ModelArray memory requirements were low even when analyzing hundreds of participants, and only had minimal change when the number of participants increased. This scalability facilitates fixel-wise statistical analyses of large-scale data resources.

### Extensibility to diverse statistical models

Brain changes across the lifespan are often nonlinear. One of the most-widely used statistical models to capture both linear and nonlinear effects is the GAM. GAMs use smooth functions to flexibly model linear and nonlinear effects; these smooth functions can be penalized to avoid over-fitting. The incorporation of GAMs in ModelArray represents an advance over existing tools, which at present only support the GLM. However, it should be noted that because ModelArray is built within R, it has the potential to leverage the myriad of statistical models that R provides. Indeed, additional statistical models can be added to ModelArray using the same workflow described in the developer documentation (https://pennlinc.github.io/ModelArray/articles/doc_for_developer.html). This extensibility will allow for ongoing enhancements – by both the original developers and the broader community – to extend ModelArray’s functionality to a wide variety of statistical models.

### Limitations and future directions

Several limitations of ModelArray should be noted. First, ModelArray is configured to only analyze fixel data. Moving forward, it may be generalized to allow for analyses of other imaging data types such as voxel (NIfTI) and surface (CIFTI) data. Such extensions could leverage ModelArray’s modular I/O interface, which would only require additional companion converters (i.e., ConVoxel instead of ConFixel). Second, ModelArray does not incorporate information of fixel-fixel connectivity (in contrast to CFE with MRtrix), which limits the ability of ModelArray to conduct cluster-wise statistical inference. However, the control of multiple comparisons using methods such as FDR is commonly used in large-scale studies and is currently implemented in ModelArray. Third and finally, ModelArray requires installation in R and depends on other R packages.

### Conclusion

ModelArray is a scalable R package for fixel-wise statistical analysis. It reduces memory requirements and offers both linear and nonlinear modeling with substantial extensibility. Taken together, ModelArray facilitates the statistical analysis of fixel data in large-scale dMRI datasets.

## ACKNOWLEDGEMENTS

This study was supported by grants from the National Institutes of Health: R01MH112847 (R.T.S., T.D.S.), R01MH120482 (T.D.S.), R37MH125829 (T.D.S.), R01EB022573 (T.D.S.), R01MH113550 (T.D.S.), R01MH123550 (R.T.S.), RF1MH116920 (T.D.S.), 5T32DC000038 (S.L.M.), K99MH127293 (B.L.), R01MH119185 (D.R.R), R01MH120174 (D.R.R.). V.J.S. was supported by a National Science Foundation Graduate Research Fellowship (DGE-1845298). Additional support was provided by the AE Foundation and the Penn/CHOP Lifespan Brain Institute.

## REFERENCES

Alexander, L. M., Escalera, J., Ai, L., Andreotti, C., Febre, K., Mangone, A., Vega-Potler, N., Langer, N., Alexander, A., Kovacs, M., Litke, S., O’Hagan, B., Andersen, J., Bronstein, B., Bui, A., Bushey, M., Butler, H., Castagna, V., Camacho, N.,… Milham, M. P. (2017). An open resource for transdiagnostic research in pediatric mental health and learning disorders. Scientific Data, 4(1), 170181. https://doi.org/10.1038/sdata.2017.181

Basser, P. J., & Pierpaoli, C. (1996). Microstructural and Physiological Features of Tissues Elucidated by Quantitative-Diffusion-Tensor MRI. Journal of Magnetic Resonance, Series B, 111(3), 209–219. https://doi.org/10.1006/jmrb.1996.0086

Bethlehem, R. A. I., Seidlitz, J., White, S. R., Vogel, J. W., Anderson, K. M., Adamson, C., Adler, S., Alexopoulos, G. S., Anagnostou, E., Areces-Gonzalez, A., Astle, D. E., Auyeung, B., Ayub, M., Bae, J., Ball, G., Baron-Cohen, S., Beare, R., Bedford, S. A., Benegal, V.,… Alexander-Bloch, A. F. (2022). Brain charts for the human lifespan. Nature, 1–11. https://doi.org/10.1038/s41586-022-04554-y

Cieslak, M., Cook, P. A., He, X., Yeh, F.-C., Dhollander, T., Adebimpe, A., Aguirre, G. K., Bassett, D. S., Betzel, R. F., Bourque, J., Cabral, L. M., Davatzikos, C., Detre, J. A., Earl, E., Elliott, M. A., Fadnavis, S., Fair, D. A., Foran, W., Fotiadis, P.,… Satterthwaite, T. D. (2021). QSIPrep: An integrative platform for preprocessing and reconstructing diffusion MRI data. Nature Methods, 18(7), 775–778. https://doi.org/10.1038/s41592-021-01185-5

Dhollander, T., Clemente, A., Singh, M., Boonstra, F., Civier, O., Duque, J. D., Egorova, N., Enticott, P., Fuelscher, I., Gajamange, S., Genc, S., Gottlieb, E., Hyde, C., Imms, P., Kelly, C., Kirkovski, M., Kolbe, S., Liang, X., Malhotra, A.,… Caeyenberghs, K. (2021). Fixel-based Analysis of Diffusion MRI: Methods, Applications, Challenges and Opportunities. NeuroImage, 241, 118417. https://doi.org/10.1016/j.neuroimage.2021.118417

Jeurissen, B., Leemans, A., Tournier, J. D., Jones, D. K., & Sijbers, J. (2013). Investigating the prevalence of complex fiber configurations in white matter tissue with diffusion magnetic resonance imaging. Human Brain Mapping, 34(11), 2747–2766. https://doi.org/10.1002/hbm.22099

Lebel, C., Gee, M., Camicioli, R., Wieler, M., Martin, W., & Beaulieu, C. (2012). Diffusion tensor imaging of white matter tract evolution over the lifespan. NeuroImage, 60(1), 340–352. https://doi.org/10.1016/j.neuroimage.2011.11.094

Marek, S., Tervo-Clemmens, B., Calabro, F. J., Montez, D. F., Kay, B. P., Hatoum, A. S., Donohue, M. R., Foran, W., Miller, R. L., Hendrickson, T. J., Malone, S. M., Kandala, S., Feczko, E., Miranda-Dominguez, O., Graham, A. M., Earl, E. A., Perrone, A. J., Cordova, M., Doyle, O.,… Dosenbach, N. U. F. (2022). Reproducible brain-wide association studies require thousands of individuals. Nature, 1–7. https://doi.org/10.1038/s41586-022-04492-9

Pagès, H. (2021). HDF5Array: HDF5 backend for DelayedArray objects (R package version 1.20.0) [Computer software]. https://bioconductor.org/packages/HDF5Array

Pagès, H., Hickey, P., & Lun, A. (2021). DelayedArray: A unified framework for working transparently with on-disk and in-memory array-like datasets. (R package version 0.18.0) [Computer software]. https://bioconductor.org/packages/DelayedArray

Pines, A. R., Larsen, B., Cui, Z., Sydnor, V. J., Bertolero, M. A., Adebimpe, A., Alexander-Bloch, A. F., Davatzikos, C., Fair, D. A., Gur, R. C., Gur, R. E., Li, H., Milham, M. P., Moore, T. M., Murtha, K., Parkes, L., Thompson-Schill, S. L., Shanmugan, S., Shinohara, R. T.,… Satterthwaite, T. D. (2022). Dissociable multi-scale patterns of development in personalized brain networks. Nature Communications, 13(1), 2647. https://doi.org/10.1038/s41467-022-30244-4

R Core Team. (2021). R: A Language and Environment for Statistical Computing (R version 4.1.2) [Computer software]. R Foundation for Statistical Computing. https://www.R-project.org

Raffelt, D. A., Smith, R. E., Ridgway, G. R., Tournier, J. D., Vaughan, D. N., Rose, S., Henderson, R., & Connelly, A. (2015). Connectivity-based fixel enhancement: Whole-brain statistical analysis of diffusion MRI measures in the presence of crossing fibres. NeuroImage, 117, 40–55. https://doi.org/10.1016/j.neuroimage.2015.05.039

Raffelt, D. A., Tournier, J. D., Smith, R. E., Vaughan, D. N., Jackson, G., Ridgway, G. R., & Connelly, A. (2017). Investigating white matter fibre density and morphology using fixel-based analysis. NeuroImage, 144(Pt A), 58–73. https://doi.org/10.1016/j.neuroimage.2016.09.029

Roalf, D. R., Quarmley, M., Elliott, M. A., Satterthwaite, T. D., Vandekar, S. N., Ruparel, K., Gennatas, E. D., Calkins, M. E., Moore, T. M., Hopson, R., Prabhakaran, K., Jackson, C. T., Verma, R., Hakonarson, H., Gur, R. C., & Gur, R. E. (2016). The impact of quality assurance assessment on diffusion tensor imaging outcomes in a large-scale population-based cohort. NeuroImage, 125, 903–919. https://doi.org/10.1016/j.neuroimage.2015.10.068

Satterthwaite, T. D., Elliott, M. A., Ruparel, K., Loughead, J., Prabhakaran, K., Calkins, M. E., Hopson, R., Jackson, C., Keefe, J., Riley, M., Mentch, F. D., Sleiman, P., Verma, R., Davatzikos, C., Hakonarson, H., Gur, R. C., & Gur, R. E. (2014). Neuroimaging of the Philadelphia neurodevelopmental cohort. NeuroImage, 86, 544–553. https://doi.org/10.1016/j.neuroimage.2013.07.064

Schilling, K. G., Daducci, A., Maier-Hein, K., Poupon, C., Houde, J.-C., Nath, V., Anderson, A. W., Landman, B. A. & Descoteaux, M. (2018). Challenges in diffusion MRI tractography – Lessons learned from international benchmark competitions. Magnetic Resonance Imaging, 57, 194–209. https://doi.org/10.1016/j.mri.2018.11.014

Tournier, J.-D., Calamante, F., & Connelly, A. (2007). Robust determination of the fibre orientation distribution in diffusion MRI: Non-negativity constrained super-resolved spherical deconvolution. NeuroImage, 55(4), 1459–1472. https://doi.org/10.1016/j.neuroimage.2007.02.016

Tournier, J.-D., Smith, R., Raffelt, D., Tabbara, R., Dhollander, T., Pietsch, M., Christiaens, D., Jeurissen, B., Yeh, C.-H., & Connelly, A. (2019). MRtrix3: A fast, flexible and open software framework for medical image processing and visualisation. Neuroimage, 202, 116137. https://doi.org/10.1016/j.neuroImage.2019.116137

Van Essen, D. C., Smith, S. M., Barch, D. M., Behrens, T. E. J., Yacoub, E., Ugurbil, K., & Consortium, for the W.-M. H. (2013). The WU-Minn Human Connectome Project: An overview. Neuroimage, 80, 62–79. https://doi.org/10.1016/j.neuroImage.2013.05.041

Wood, S. N. (2001). mgcv: GAMs and generalized ridge regression for R. R News, 1/2, 20–25.

Wood, S. N. (2004). Stable and efficient multiple smoothing parameter estimation for generalized additive models. Journal of the American Statistical Association, 99(467), 673–686. https://doi.org/10.1198/016214504000000980

Yeh, F.-C., Verstynen, T. D., Wang, Y., Fernández-Miranda, J. C. & Tseng, W.-Y. I. (2013). Deterministic Diffusion Fiber Tracking Improved by Quantitative Anisotropy. PLoS ONE, 8(11), e80713. https://doi.org/10.1371/journal.pone.0080713

